# Alignment-Free Viral Sequence Classification at Scale

**DOI:** 10.1101/2024.12.10.627186

**Authors:** Daniel J. van Zyl, Marcel Dunaiski, Houriiyah Tegally, Cheryl Baxter, The INFORM Africa research study group, Tulio de Oliveira, Joicymara S. Xavier

## Abstract

**Background:** The rapid increase in nucleotide sequence data generated by next-generation sequencing (NGS) technologies demands efficient computational tools for sequence comparison. Alignment-based methods, such as BLAST, are increasingly overwhelmed by the scale of contemporary datasets due to their high computational demands for classification. This study evaluates alignment-free (AF) methods as scalable and rapid alternatives for viral sequence classification, focusing on identifying techniques that maintain high accuracy and efficiency when applied to extremely large datasets.

**Results:** We employed six established AF techniques to extract feature vectors from viral genomes, which were subsequently used to train Random Forest classifiers. Our primary dataset comprises 297,186 SARS-CoV-2 nucleotide sequences, categorized into 3502 distinct lineages. Furthermore, we validated our models using dengue and HIV sequences to demonstrate robustness across different viral datasets. Our AF classifiers achieved 97.8% accuracy on the SARS-CoV-2 test set, and 99.8% and 89.1% accuracy on dengue and HIV test sets, respectively.

**Conclusion:** Despite the high-class dimensionality, we show that word-based AF methods effectively represent viral sequences. Our study highlights the practical advantages of AF techniques, including significantly faster processing compared to alignment-based methods and the ability to classify sequences using modest computational resources.

## Introduction

Amidst the plethora of groundbreaking contributions made to the field of molecular biology in the 1980’s and 1990’s, perhaps one of the most significant was an algorithm designed by two mathematicians [1]. In 1990, Karlin and Altschul proposed a means to approximate the similarity between two DNA/RNA sequences and to do so an order of magnitude faster than existing alternatives [2]. The algorithm would then become the statistical foundation of BLAST [3], the Basic Local Alignment Search Tool, one of the most successful alignment-based comparison tools ever created [1, 4].

Beyond its direct impact on the field, the success of BLAST also marked a significant developmental shift toward the refinement of probabilistic models for alignment-based sequence analysis, along with the development and application of advanced string matching algorithms [4]. Regardless of implementation, alignment-based sequence comparison tools seek to identify regions of similarity between sequences through the matching of nucleotide bases that occur in the same order between the given sequences [1]. These methods, while fundamental to genomic research, require considerable computational resources, noting that there is an exponential increase in possible alignments with increasing sequence length [1, 5].

The introduction of next-generation sequencing (NGS) technologies has reshaped the landscape of computational biology. In 1990, when BLAST was introduced, there were fewer than 50 million nucleotide bases publicly available. Today, a single sequencing instrument has the capability to generate more than 1 trillion bases per run [6]. The sheer volume of data produced by NGS is rapidly exceeding the capabilities of analytics tools, largely due to the intensive computational requirements of the multiple alignment process [1]. Hence, contemporary research is increasingly turning to alignment-free (AF) methods as viable alternatives for sequence comparison [5, 7–9].

AF sequence comparison techniques have undergone extensive research, development, and benchmarking over several decades [9, 10]. Despite demonstrating remarkable performance on general biological sequence datasets in numerous studies, the widespread adoption of AF sequence classification remains limited. Two primary reasons may contribute to this hesitation. Firstly, datasets commonly utilized to evaluate AF techniques often contain trivial cases, featuring only a small number of distinct species [10]. Secondly, we contend that the choice of datasets may not align with real-time scenarios where the rapid nature of AF sequence comparison could provide significant benefits.

Monitoring viral pathogen strains in near real time is a prominent application of AF sequence comparison methods, yet little research has assessed their ability to accurately represent viral sequences. This is particularly prevalent, as alignment-based tools depend on sequence collinearity, the preservation of homologous nucleotide order across genomes, an assumption frequently violated in viral genomes due to high mutation rates and frequent recombination events [1].

In this study, we conducted a comprehensive evaluation of AF methods for classifying viral sequences at scale. We used six established AF sequence comparison techniques to extract representative feature vectors from viral genomes, serving as input for Random Forest classifiers. We applied our AF models to a large-scale dataset of 297,186 SARS-CoV-2 nucleotide sequences, assessing their ability to classify sequences into 3502 distinct lineages. Given the size of the dataset, the considerable number of target classes, and the high similarity understood among strains of the same virus, this experiment represents, to the best of our knowledge, one of the most intensive evaluations of AF sequence classification in the literature. Furthermore, we validated the effectiveness of our models using moderately sized datasets of HIV and dengue sequences.

We provide the source code in the form of a library of AF feature extraction methods for custom model development along with a command-line tool to classify HIV, dengue, and SARS-CoV-2 sequences^1^. Additionally, the tool is adaptable to other viral pathogens and includes straightforward training infrastructure.

## Backround

AF approaches can fall under a number of different methodologies, including those relying on the frequencies of subsequences of a specific length (oligomeric/word-based methods) [7, 8, 11, 12], those rooted in information theory [13–16], those based on the length of matching words or common substrings [17–19], and other unique methods that include chaos game representations [20] and digital signal processing [21, 22]. Regardless of the underlying implementation, AF techniques typically transform biological sequences into numeric feature vectors that are used to compute pairwise dissimilarity scores to construct phylogenetic models.

However, AF comparison tools can also be seen as a means of direct feature extraction for the purposes of machine learning. In this context, feature extraction refers to the transformation of biological sequences into vectors that numerically describe the characteristics of the original sequences in a way that maximizes information gain while minimizing potential noise [23].

Zielezinski et al. [10] highlight several challenges that hinder the widespread adoption of AF methodologies. They point out the lack of standardization in evaluation strategies, benchmark datasets, and test criteria. More critically, they argue that new methods are frequently evaluated using small, non-representative datasets chosen by their authors, and are often only validated against a limited selection of alternative AF approaches. To address these limitations, Zielezinski et al. developed AFproject, an online service designed to benchmark AF tools in various sequence analysis scenarios, including protein sequence classification, gene tree inference, regulatory sequence identification, genome-based phylogenetics, and horizontal gene transfer. With regard to the use of machine learning methods, Bonidia et al. [9] analyzed AF feature extraction approaches for biological sequence classification, with the objective of evaluating the ability of mathematical features to generalize across different long noncoding RNA lncRNA classification tasks.

Neither Zielezinski et al. nor Bonidia et al. evaluated AF tools on viral datasets. As part of a novel machine learning method CASTOR-KRFE, Lebatteux et al. [5] evaluated their method on a dozen diverse virus datasets, covering the seven major virus groups. The datasets included influenza virus, Ebola virus, human immunodeficiency virus 1, hepatitis C virus, hepatitis B virus, and human papillomavirus. However, their largest dataset consists of only 1352 samples with only 28 classification targets. Lebatteux et al. did not evaluate alternative feature extraction methodologies.

Regarding SARS-CoV-2, the largest source of virus sequence data currently available, Muhammad et al. [24] implemented two boosting algorithms, eXtreme Gradient Boosting (XGBoost) and Light Gradient Boosting Machine (LGBM), to classify sequences of SARS-CoV-2. However, their study only included variants of concern: alpha, beta, gamma, delta, and omicron (VOC); as well as variants of interest (VOI).

One of the largest studies of machine learning to date to classify SARS-CoV-2 sequences is that of Lebatteux et al. [25], who compared the machine learning tools KEVOLVE and CASTOR-KRFE with statistical tools to identify discriminative motifs in unaligned sequence sets to classify SARS-CoV-2 variants. They constructed a comprehensive dataset of 334,956 SARS-CoV-2 genomes, but only included genomes with ambiguous nucleotides less than 1% and only covered 10 major variants with more than 100 samples each.

Cacciabue et al. [26] introduced Covidex, an open source and alignment-free machine learning tool designed for subtyping SARS-CoV-2 based on k-mer frequency profiles, which are used as input features for Random Forest classifiers. The tool achieved 96.56% accuracy in distinguishing among 1,437 Pango lineages of SARS-CoV-2 as of late 2021. Despite its success, the study did not explore alternative feature extraction methods or extend its classification framework to other viruses.

## Results and Discussion

Table 1 lists the key performance results of the selected feature extraction techniques on the dengue, HIV and SARS-Cov-2 hold out test sets across three key metrics, accuracy, Macro F1 score and Mathew’s Correlation Coefficient (MCC).

**Table 1.**
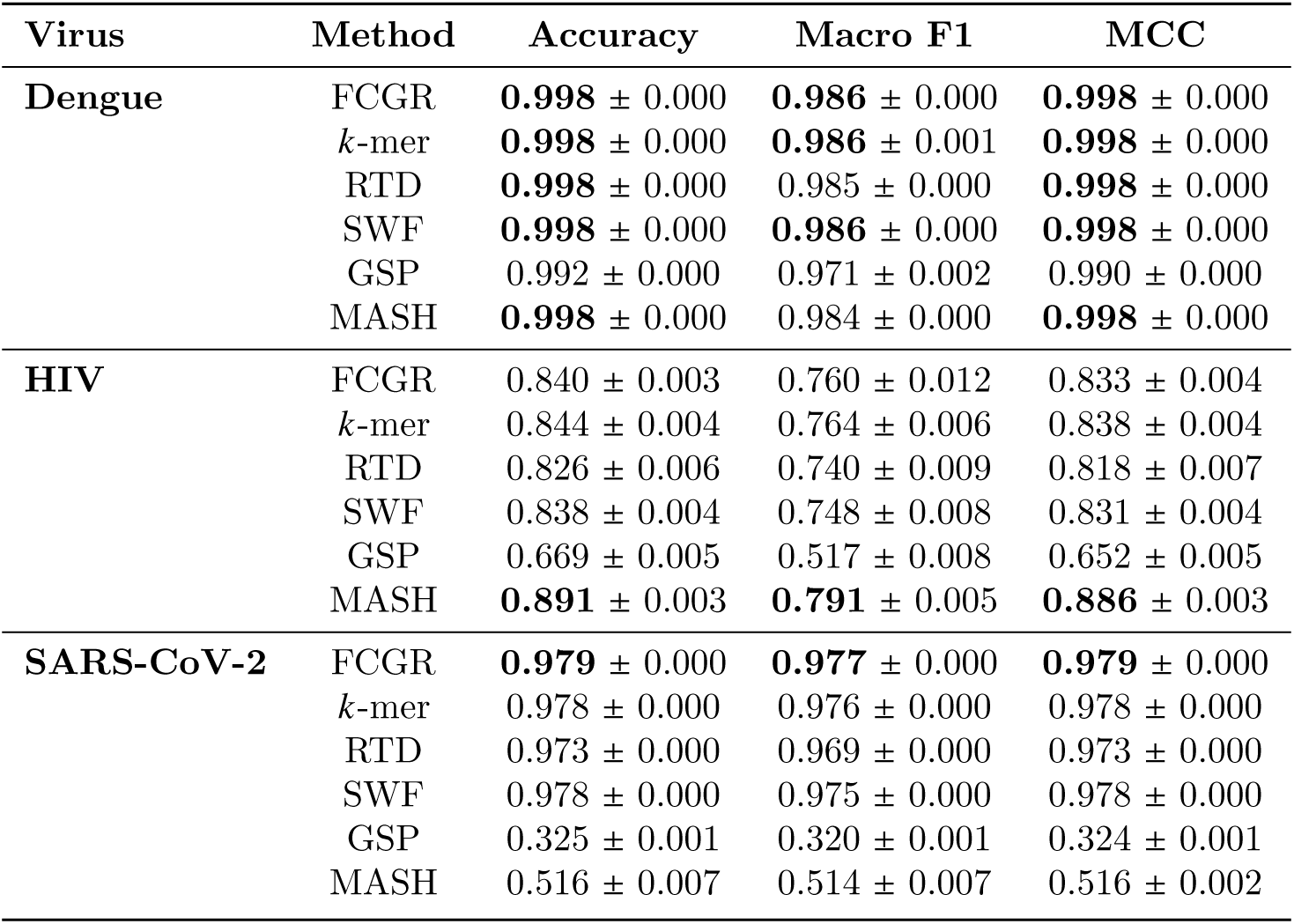
Summary of the performance results for various AF feature extraction methods applied to dengue, HIV, and SARS-CoV-2 hold-out test sets. For each method, the accuracy, Macro F1 Score, and Matthews Correlation Coefficient (MCC) are reported with their respective standard deviations.

The selected feature extraction techniques include *k*-mer counting, Frequency Chaos Game Representation (FCGR) [20], Return Time Distribution (RTD) [7], Spaced Word Frequencies (SWF) [8], Genomic Signal Processing (GSP) [21, 22] and Mash [6].

The optimal parameter configurations and strategies for handling degenerate nucleotides remained consistent for each feature extraction technique across all datasets. Consequently, the optimal configurations are summarized in Table 2. See Additional file 1 for an analysis of feature extraction parameters on the SARS-CoV-2 dataset.

**Table 2.** Optimal parameter configurations and degeneracy handling for each feature extraction method.

| AF Method | Parameters | Degenerates |
| --- | --- | --- |
| FCGR | $r = 128$ | Removed |
| *k*-mer | $k = 7$ | Removed |
| RTD | $k = 7$ | Removed |
| SWF | $k = 7$ | Removed |
| GSP | Real Mapping | Replaced |
| MASH | $k = 21$; $s = 1000$ | Removed |

### Dengue

FCGR, SWF, *k*-mers, RTD, and MASH all achieve near-perfect classification performance for dengue sequence classification. All five methods achieve an overall classification accuracy and MCC of 99.8%, with FCGR, SWF *k*-mers achieving the highest Macro F1 score of 98.6%.

The exceptionally high performance results for these five feature extraction techniques indicate a strong suitability for AF sequence classification in the context of dengue classification at the genotypic level. However, the slight discrepancy between the accuracy scores and the Macro F1 scores indicates a slight bias toward the majority classes.

The outlying feature extraction technique, GSP, still performs reasonably well with a classification accuracy greater than 99%. In previous work, Randhawa et al. [21] demonstrated that signal processing feature extraction techniques could achieve perfect accuracy at the Serotypic level. Thus, our findings suggest that when the classification task is extended to the genotypic level, the performance of GSP begins to degrade slightly.

### HIV

Among the evaluated methods, Mash stands out with the highest accuracy (0.891), Macro F1 Score (0.793), and MCC (0.886). In general, the classification performance of all feature extraction techniques was significantly lower in the case of HIV classification than in the case of dengue classification. Similarly to the dengue dataset, we found that the respective models all achieved lower Macro F1 scores than accuracy scores, however, in the case of HIV, this discrepancy was larger, indicating that the models struggled to classify minority classes.

Following Mash, the word-based methods FCGR, *k*-mers, RTD and SWF all achieve similarly high performance scores of above 80% accuracy and MCC and 70% Macro F1. However, GSP performed significantly worse than the other methods, achieving only a classification accuracy of 55. 8% in the HIV dataset.

### SARS-CoV-2

Among the methods evaluated, FCGR proved to be the most effective, achieving the highest accuracy (0.979), Macro F1 Score (0.977), and MCC (0.979). Similarly, *k*-mer RTD (0.978) and SWF (0.974) also achieve high accuracy, with the top-performing methods all utilizing word counting techniques. In contrast, GSP achieves notably poor performance, with an average accuracy of only 0.325, making it the least effective method by a significant margin. All models produce similar and consistent results across all evaluation metrics, indicating that all models achieve balanced performance across different classes independent of class size.

Table 3 compares Macro F1 scores between training, validation, and test sets. We consider Macro F1 to be the most robust performance metric, as it more effectively accounts for class imbalances and bias. All models exhibit a higher (near perfect) performance in the training set compared to the validation and test sets. This discrepancy is most pronounced with GSP, which achieves perfect accuracy on the training set but relatively poor accuracy on the validation and test sets. This pattern typically indicates overfitting, where the models learn the noise in the training data, reducing their generalization ability. However, for this application, increasing the complexity of the Random Forest models results in the best generalization performance, although the improvement in generalization did not match the rate of improvement in training performance with increasing model depth. The validation and testing results are strongly correlated across all models, indicating consistent performance on unseen data and confirming the reliability of the validation results as predictors of testing performance.

**Table 3.** Comparison of Macro F1 performance across the train, validation and test sets for the SARS-CoV-2 dataset.

| Model | Set |  |  |
| --- | --- | --- | --- |
|  | Train | Validation | Test |
| SWF | 0.9999 | 0.9756 | 0.9754 |
| *k*-mer | 0.9999 | 0.9766 | 0.9755 |
| MASH | 0.9981 | 0.5133 | 0.5144 |
| FCGR | 0.9999 | 0.9773 | 0.9768 |
| GSP | 1.0000 | 0.3189 | 0.3149 |
| RTD | 0.9999 | 0.9697 | 0.9687 |

### Class-wise Classification Performance

We analyzed the class-wise accuracy of each model to determine whether the models perform uniformly across all classes or whether they excel or underperform for certain sets of classes. Figure 1 gives the class-wise accuracy results for the different models. The SARS-CoV-2 lineages (classes) are ordered in descending order of the average classification performance across all models. The figure demonstrates the overall classification behavior of the models and the rate at which performance degrades when models face lineages that are more challenging to classify correctly.

**Fig. 1.**
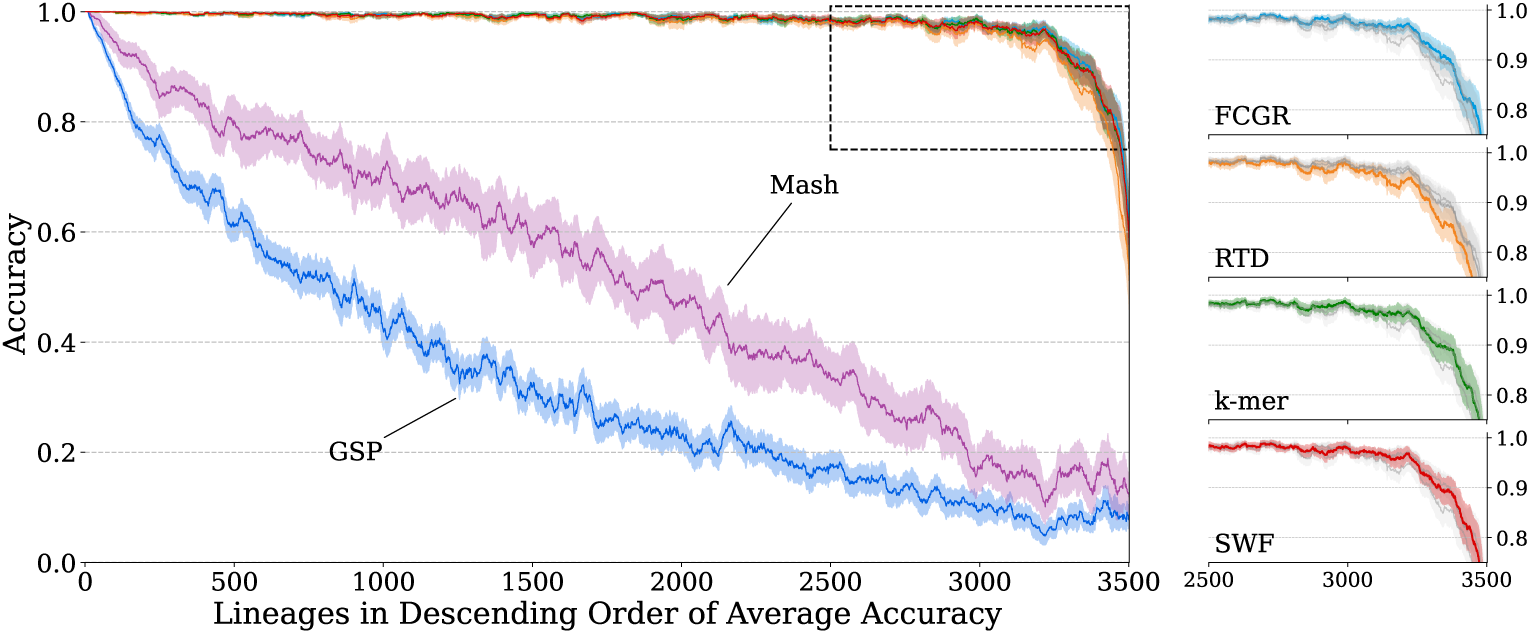
The SARS-CoV-2 testing accuracy results of each AF feature extraction method on a classwise basis. The findings provide insights into the distribution of model performance across classes. Classes are ordered in descending order of average classification accuracy across all models. The standard deviations of the accuracy for each model are also depicted. For the purposes of visual clarity, the values depicted have been smoothed using a sliding window of 50 classes. On the right-hand side, we provide isolated views of the top performing models.

Our findings again indicate that word-based models achieve the best results, while also displaying remarkably similar behavior. These models achieved near-perfect accuracy for the vast majority of the 3502 classes in the test set, with a sharp decline in performance observed in only a few classes. Furthermore, in the vast majority of cases in which these models achieve perfect accuracy, they did so with zero deviation. The word-based models exhibit a steep decline in performance in only around 200 classes. In contrast, GSP showed an immediate and notable decline in performance, while Mash displayed an almost linear decline in class-wise accuracy. Both GSP and Mash also demonstrated a higher degree of performance deviation throughout.

### Factors Influencing Classification Performance

Given that even the best models exhibit notably poor performance in a small minority of classes, we investigated potential reasons for this behavior. One possible challenge identified in the classification of SARS-CoV-2 sequences is the presence of recombinants, lineages resulting from the combination of genetic material from different lineages of the virus, leading to new hybrid sequences. This recombination process can introduce inherent noise that can complicate the classification task for machine learning models. To assess the impact of recombinants on model performance, we compared the class-wise accuracy of the models on nonrecombinant sequences and on recombinants, as shown in a set of split violin plots in Figure 2.

**Fig. 2.**
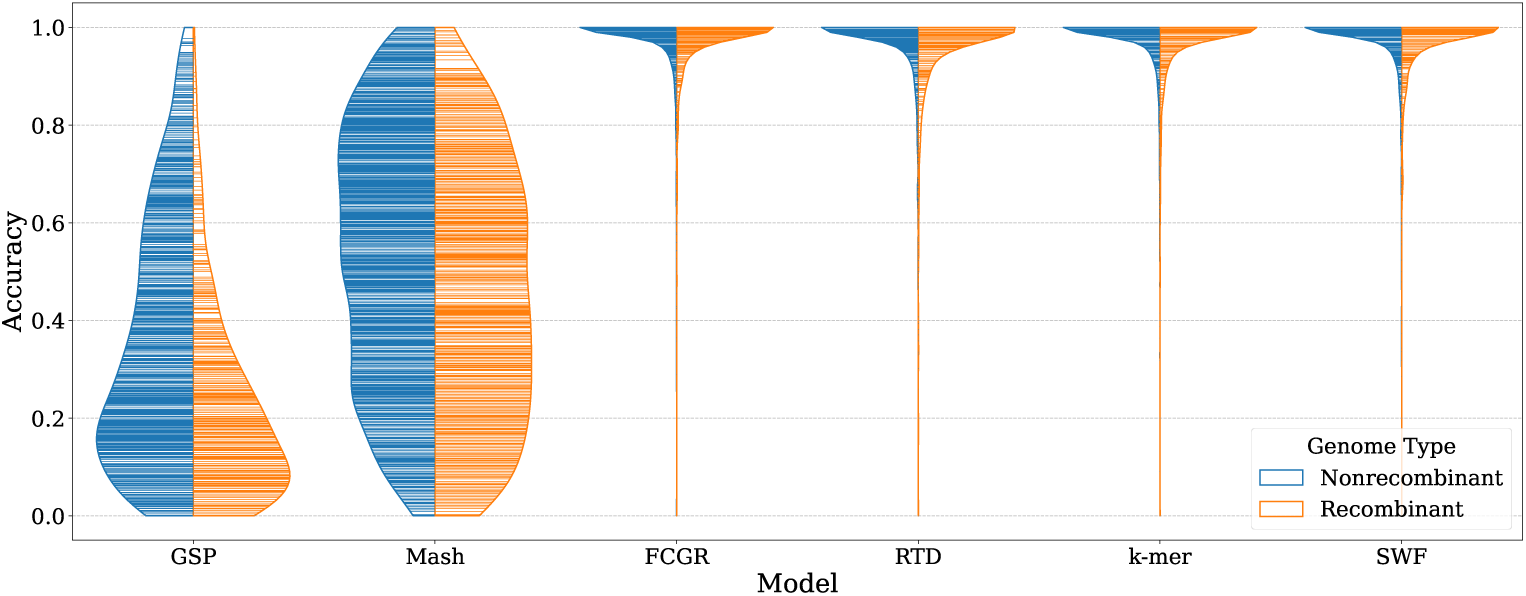
Comparison of the performance of each AF feature extraction model on nonrecombinant genomes (blue) and recombinant (orange) genomes. The horizontal lines in each violin plot indicate the models’ achieved accuracy for different individual classes, while the width of the violin plots represents the density of samples at different accuracy levels.

The results indicate that recombinants are indeed more challenging to classify across all models. The performance drop is most pronounced for Mash and GSP, which struggle significantly with recombinant sequences. In contrast, word-based models show only a slight decrease in performance.

We further investigated three additional factors that could potentially influence model performance: the number of available training samples, the evolutionary depth of the sequences, and the number of direct descendants, also known as sublineages, that each lineage possesses.

Figure 3 presents a grid of hexbin correlation plots, with each column corresponding to one of these three variables.

**Fig. 3.**
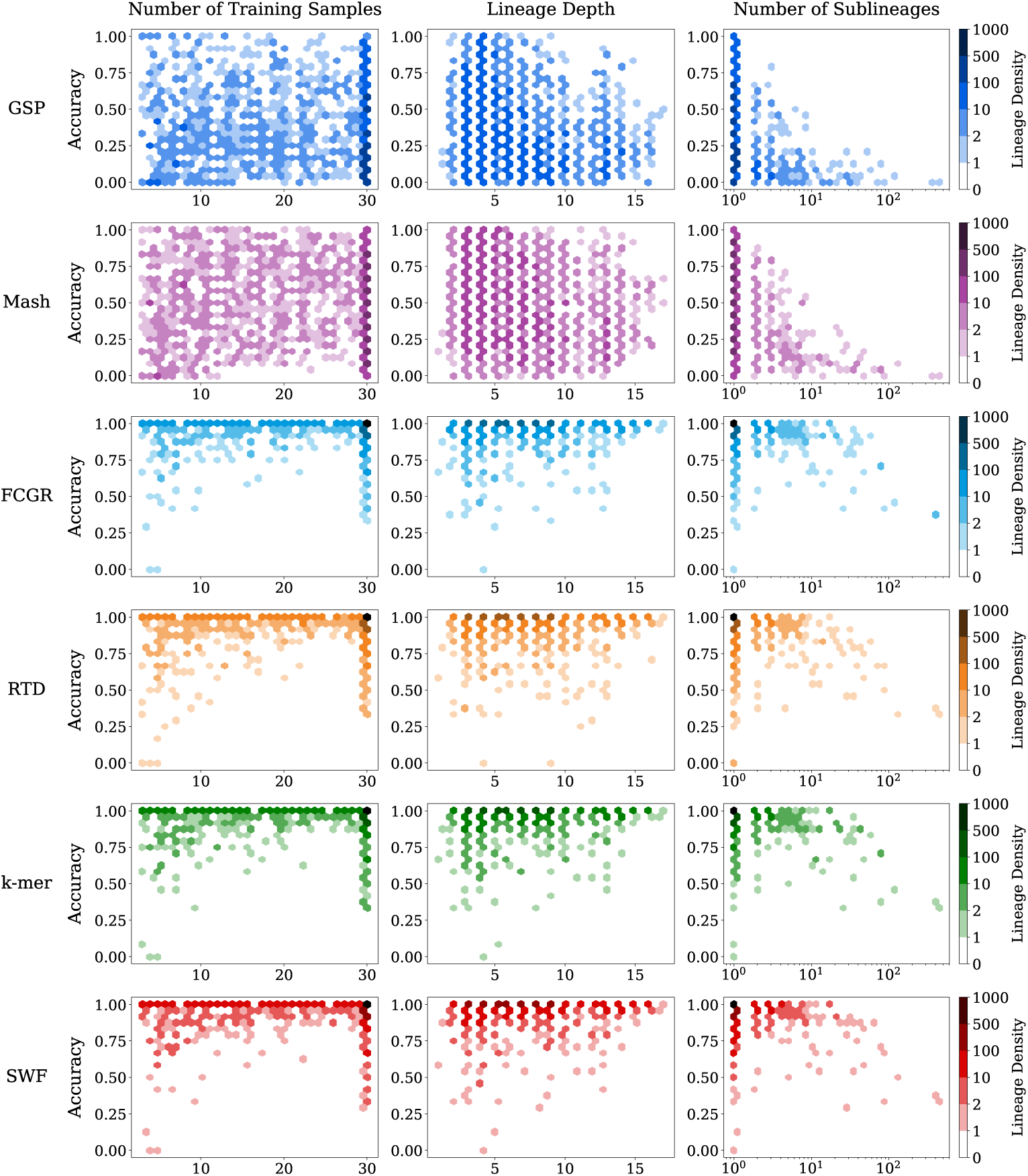
This figure depicts the interaction between classification accuracy and the number of training samples for each class (left), the depth of each lineage (middle), and the number of direct descendants of each lineage (right) for each SARS-CoV-2 model in the form of hexbin plots. We only show depth and number of descendant interactions for nonrecombinant, nonrecombinant sequences.

With regard to the number of training samples available, it is evident that word-based models exhibit a noticeable decrease in classification accuracy when fewer training samples are available. Lineages with fewer than 10 training samples demonstrated a proportionally higher likelihood of suboptimal classification accuracy. However, a significant proportion of lineages with suboptimal accuracy also had the maximum available training sample size of 30. Although this is consistent with the general observation that most lineages have 30 training samples, it suggests that factors beyond the quantity of training samples also contribute to reduced classification performance. For Mash and GSP, less notable interactions are evident.

Across all models, no significant relationship was observed between lineage depth and model performance. Although most lineages with suboptimal classification accuracy had an evolutionary depth of less than 10, this trend reflects the higher overall representation of such lineages in the dataset rather than a direct influence of depth on performance.

Lastly, all models exhibited a significant decline in classification accuracy as the number of sublineages increased. This performance degradation was more pronounced than the impact of limited training samples, and all models consistently fell below perfect accuracy once a universal sublineage threshold was exceeded.

As a final evaluation of our models, we investigated the performance of the three best performing models (FCGR, *k*-mers, and SWF) on the most frequently occurring lineages during the SARS-CoV-2 pandemic. This analysis provides a holistic view of model performance in a more tangible and recognizable set of samples.

Figure 4 shows a radar chart of classification performance in the 200 most common nonrecombinant lineages. The surrounding circular bar chart represents the evolutionary depth of each lineage, color-coded according to the clades to which the lineages belong. The findings show remarkably similar behavior across all models. Consistent with the overall results, the models perform very well in the majority of classes, with significant dips for certain lineages. Model performance does not fluctuate significantly between different clades. However, we observe significant performance drops for lineages B (one of the two original haplotypes), B.1.617.2 (Delta), B.1.1.529 (Omicron), B.1.1, BA.2 (a direct descendant of Omicron) and JN.1. These performance drops are particularly notable in some of the most critical strains of the SARS-CoV-2 virus, especially Delta and Omicron.

**Fig. 4.**
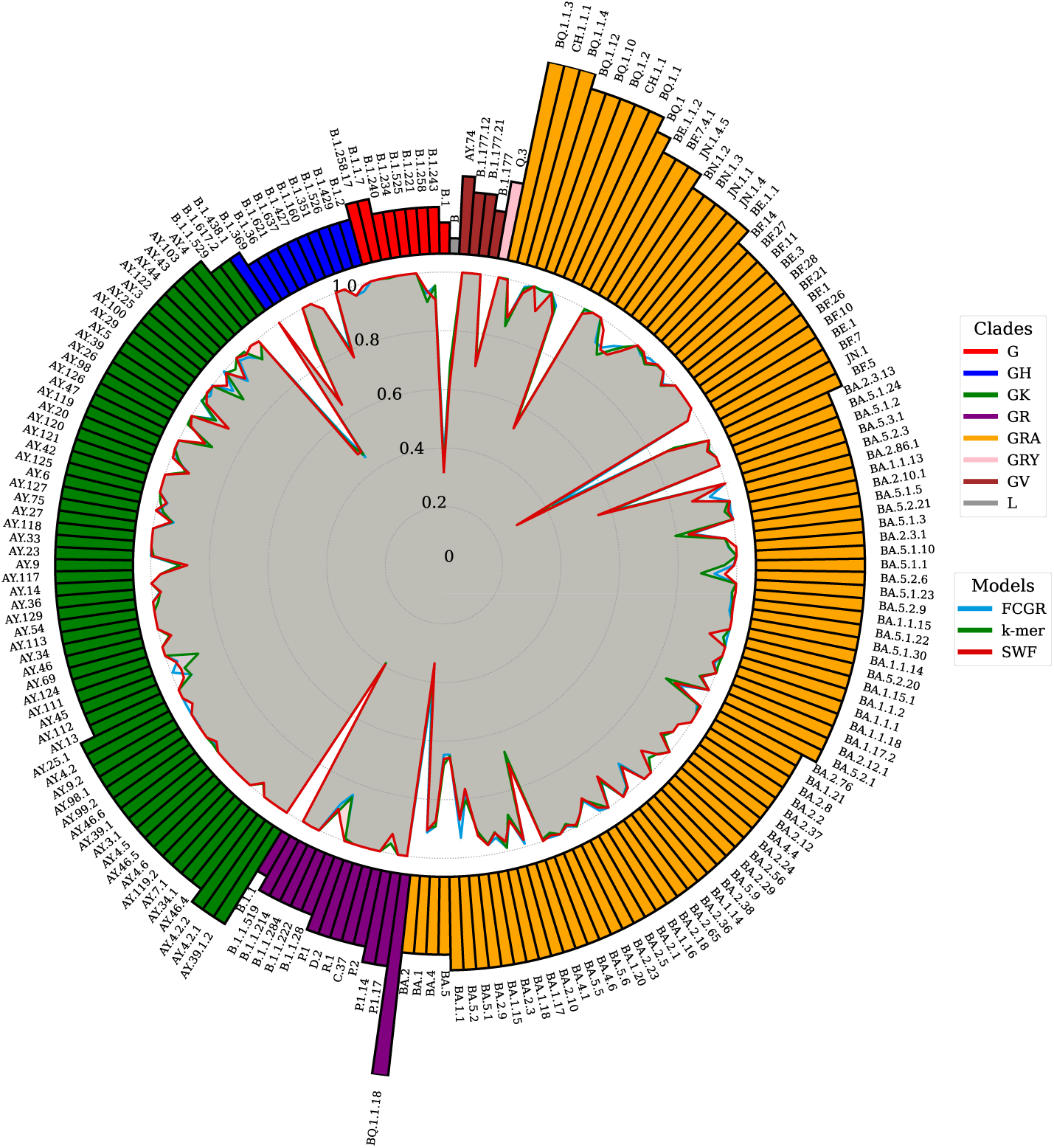
A composite figure of the class-wise classification performance of the top three performing models, FCGR,*k*-mers, and SWF on the 200 most promintent SARS-CoV-2 lineages. The inner plot consists of a radar chart, where optimal performance corresponds to observations near the perimeter of the chart. The outer figure shows a circular bar plot in which the bars correspond to the depth of the SARS-CoV-2 lineages and are colored according to the respective clades of the lineages.

## Conclusions

In this paper, we comprehensively evaluated the effectiveness of alignment-free (AF) methods for the classification of viral sequences on a large scale, focusing on both classification performance and computational efficiency. We used six established AF techniques to extract feature vectors from viral genomes and trained Random Forest classifiers on these features. We experimented with various parameters for AF techniques and evaluated the effects of removing versus replacing degenerate nucleotides. The findings from our primary dataset, consisting of 297,186 SARS-CoV-2 nucleotide sequences categorized into 3,502 distinct lineages, demonstrate that AF methods can achieve high classification accuracy and efficiency even with large-scale datasets. We also evaluated class-wise accuracy based on the number of sublineages, training samples, and the depth of each class to determine the factors that influence classification difficulty.

Despite the high dimensionality, we found that word-based AF methods could effectively represent the SARS-CoV-2 sequences, achieving classification accuracies close to 98% on the test set. Furthermore, we validated our models using moderately sized HIV (4,177 samples) and Dengue (18,970 samples) datasets to ensure their robustness across different viral types.

Our findings further contribute to the practical advantages of AF sequence classification. AF viral sequence classification is considerably faster than alignment-based techniques. Moreover, the ability to train models and classify viral sequences using only modest computational resources, without reliance on cloud infrastructure, underscores the accessibility and scalability of these methods for researchers worldwide.

The application of AF sequence classification holds significant potential for future research, such as the ability to directly classify unassembled sequences on a large scale.

## Methods

We evaluated six different AF techniques, three based on word frequencies, one based on chaos theory, one based on digital signal processing, and one based on frequency of word matches. We selected this subset of AF approaches in an attempt to evaluate classification performance and computational efficiency across a diverse set of feature extraction approaches, with a focus on some of the most popular techniques. The selected AF techniques were also chosen for their ability to be directly implemented without relying on third-party software. We also investigated the use of average common substrings (ACS) but concluded it computationally infeasible for computing distance matrices for large-scale datasets.

### Feature Extraction Methods

Below is a brief overview of each selected feature extraction technique. For a more detailed breakdown of the workings of the selected techniques, see Additional file 2.

- *k*-mer counting involves breaking genomic sequences into overlapping subsequences of length *k*, called *k*-mers, and constructing a feature vector by counting the occurrences of each possible *k*-mer, with larger *k* values offering greater resolution for distinguishing similar sequences but at higher computational costs.
- Return Time Distribution (RTD) [7] provides an alternative to direct *k*-mer frequency by measuring the mean and standard deviation of the intervals between occurrences of each *k*-mer, creating a feature vector twice the size of the total possible *k*-mers and offering a distinctive perspective on sequence structure.
- Spaced Word Frequencies (SWF) [8] refine word frequency analysis by using patterns that alternate between required matches and flexible positions, focusing only on matching positions to account for sequence variations or mutations, and reducing statistical biases from consecutive matches in contiguous word approaches.
- Mash [6] adapts the MinHash technique to genomic datasets by hashing and sorting *k*-mers, creating a compact “sketch” of the smallest hash values for efficient similarity estimation. The Mash distance, derived from the Jaccard index between sketches and modeled as a Poisson process, quantifies sequence similarity based on shared hashes.
- The Frequency Chaos Game Representation (FCGR) [20] is a visualization technique for genomic sequences that maps nucleotides to positions within a unit square, starting from the center and iteratively plotting points halfway toward corners representing bases A, C, G, and T, producing a distinctive pattern that can be analyzed as frequency distributions within grid regions.
- Genomic Signal Processing (GSP) [21, 22] applies Digital Signal Processing methods to DNA/RNA sequences by converting nucleotides into numerical values using pre-defined mappings, normalizing sequence lengths, and applying the Discrete Fourier Transform (DFT) to generate frequency domain representations. The resulting magnitude spectra can be compared between sequences using Pearson correlation dissimilarity.

### Parameters

All of our selected AF feature extraction techniques incorporate parameters that influence/impact the resulting feature vectors. AF sequence comparison techniques were not originally developed as a means for feature extraction but as a means to calculate (dis)similarity between sequences. Therefore, when using AF comparison techniques to extract feature vectors, we can group these techniques into two categories. Techniques whose feature vectors are the direct result of a numeric transformation (*k*-mers, spaced word frequences, RTD and FCGR), and techniques whose feature vectors result from a derived distance measure (Mash and GSP).

*k*-mers, RTD, and SWF similarly derived from the statistical analysis of word frequencies, share a common parameter *k*, prescribing the length of the words employed. We considered values of *k* ranging between two and seven, noting diminishing performance returns against a backdrop of exponentially increasing computational demands for higher *k* values.

The key parameters for the Spaced Word Frequencies method define the selected pattern. The weight of a pattern refers to the number of match positions in the pattern and corresponds to the length of the resulting word counts. Similarly to the work of the original paper [8], for each potential weighting *k* (the number of match positions), we generated 50 random patterns with a maximum of 30 non-match positions. We then evaluated the classification performance of each pattern on the validation set and selected the pattern with the highest macro F1 score per weighting. We considered weighting values ranging between two and seven, corresponding to the word lengths selected for the other word-based techniques.

Mash also uses a word size *k*, but the ideal range of *k* values extends beyond those of the word-based methods. We investigated word sizes of 13, 19 and 21, as suggested in the original paper [6]. The second key parameter of Mash is the sketch size, for which we compared values of 500, 1000 and 2000.

GSP comprises two key parameters, the choice of numeric mapping to use and the means of length normalization. For this study, we evaluated the mappings “Real”, “PP”, and “Just-A” since these were found to be most promising in the original study [21]. For length normalization, we utilized antisymmetric median length padding [21] over zero padding [22], noting that zero padding was more likely to result in the length of sequences over-influencing classification.

Lastly, for the Frequency Chaos Game Representation, we flattened the generated images flattened into one-dimensional feature vectors. The tuning parameter *r* controls the image resolution. We considered *r* values 32, 64, and 128.

Table 4 provides an overview of the selected feature extraction techniques and their relevant parameters.

**Table 4.** Summary of feature extraction parameters. *k* refers to word length, *s* to sketch size, *r* to resolution, *l* to number of non-match positions, and *n* to numeric mapping.

| Method | Considered Parameters |
| --- | --- |
| $k$-mer | $k \in [2, 3, \dots, 7]$ |
| Return Time Distribution (RTD) [7] | $k \in [2, 3, \dots, 7]$ |
| Spaced Word Frequency (SWF) [8] | $k \in [2, 3, \dots, 7]; l = 30$ |
| Mash [6] | $s \in \{500, 1000, 2000\}; k \in \{13, 19, 21\}$ |
| Frequency Chaos Game Rep. (FCGR) [20] | $r \in \{32, 64, 128\}$ |
| Genomic Signal Processing (GSP) [21, 22] | $n \in \{\text{“Real”}, \text{“PP”}, \text{“Just A”}\}$ |

Across all feature extraction techniques, an additional universal decision lies in the handling of degenerate/ambiguous nucleotides. In this study, we considered two approaches, removing these nucleotides or randomly replacing them with viable bases.

### Data

We applied the selected AF feature extraction techniques to three viral sequence datasets. For each dataset, we partitioned the data into three distinct sets: a training set, a validation set, and a testing set, with a ratio of 50:20:30, respectively. The sets were stratified by lineage to maintain relative class balance.

### Dengue

The dengue dataset, sourced from The Global Initiative on Sharing All Influenza Data (GISAID) [27], included all available sequences as of June 2024, classified into four serotypes, each comprising multiple genotypes. Although related studies typically focus solely on classifying the four serotypes [5, 21], our study performed classification at the genotypic level.

During dataset construction, we filtered out sequences labeled as low-coverage, defined as those with more than five percent ambiguous nucleotides. To maintain statistical validity, we discarded genotypes with fewer than ten observations. One significant challenge of using machine learning for viral classification is the need for an adequate sample size for training and robust evaluation. Consequently, this filtering process led to the exclusion of two genotypes, DENV4 - III and DENV3 - IV. The remaining dataset consisted of 17,424 samples, divided into 16 different genotypic classes.

### HIV

Our HIV dataset was sourced from the Los Alamos National Lab (LANL) [28] HIV sequence database. We selected all complete genome sequences with coverage greater than 95%. Similarly to the dengue dataset, we filtered out all lineages with fewer than ten total samples. To mitigate potential bias towards common lineages, we performed random undersampling, ensuring that no single lineage had more than 300 samples. The final HIV dataset comprised 4,177 samples, divided into 53 lineages.

### SARS-CoV-2

Our large scale SARS-CoV-2 dataset was sourced from GISAID. Unlike most approaches that focus solely on dominant variants, our study expanded the classification to encompass almost all SARS-CoV-2 lineages (3,502/3,565), as defined by the Pango lineage classification nomenclature at the time of this analysis. [29].

Similarly to our approach with the dengue dataset, we removed sequences labeled as low coverage, retaining those with coverage exceeding 95%. In particular, our criteria for the tolerance of ambiguous nucleotides (95 percent coverage) are less stringent compared to similar studies, which often require more than 99 percent coverage [24, 25].

Consistent with our methodology for all datasets, we excluded classes with fewer than ten total observations from our analysis. In the case of SARS-CoV-2, this led to the elimination of 63 classes. However, the remaining classes exhibited significant class imbalance. To mitigate this bias, we used random undersampling, limiting each class to a maximum of 100 training observations. This approach ensured a balanced training set, and no class showed a disproportionate representation that exceeded a 1:10 ratio compared to any other class. The final dataset comprises 297,186 samples distributed across 3,502 distinct classes.

### Model Selection and Validation

Following the construction of various feature sets, we performed model selection and validation. We limited our scope to models that would be feasible for implementation across all datasets, and thus needed to align with the computational demands of the SARS-CoV-2 dataset. Traditional classifiers like Logistic Regression and Support Vector Machines, whilst demonstrating notable performance in similar studies, were not considered for this analysis owing to their computational demands, necessitating the use of One-vs-Rest classification strategies. We also assessed the suitability of K-Nearest Neighbours (KNN) classifiers, but found that their performance was consistently poor, likely due to the high dimensionality of the feature sets.

Considering the scale of the SARS-CoV-2 dataset and its large number of lineages, our approach naturally gravitated toward the use of tree-based algorithms, known for their favorable performance-efficiency balance on tabular datasets. We preliminarily assessed several options, including Random Forests, XGBoost, LightGBM, and CatBoost, ultimately concluding that Random Forests consistently yielded more promising performance whilst simultaneously being most computationally efficient. Consequently, we focused our investigation on the use of Random Forests. Random Forests further benefit from built-in feature selection, allowing the models to cope with the high dimensionality of the viral feature sets. We treated each model as a flat classifier, opting to overlook the hierarchical class structure inherent within the dataset and treat all classes as independent.

We used the validation set to tune the hyperparameters of each model and feature extraction technique using a grid search methodology. For each Random Forest, we investigated the use of both unweighted and balanced class weighting schemes, as well as the use of gini and entropy splitting criteria. Furthermore, we allowed the forests to grow to an unlimited maximum depth, noting that this improved performance universally. All Random Forests were constructed with 100 decision tree members, noting that in all cases, we observed diminishing returns before this point.

## Supporting information

Analysis of the feature extraction parameters on the SARS-CoV-2 dataset.

Detailed description of the workings of the selected feature extraction techniques.

Supplemental table acknowledging the authors of the SARS-CoV-2 GISAID sequences.

Supplemental table acknowledging the authors of the dengue GISAID sequences.

## Declarations

### Ethics approval and consent to participate

Not applicable

### Consent for publication

Not applicable

### Availability of data and materials

The findings of this study are based on 297,186 SARS-CoV-2 sequences available on GISAID up to and accessible at http://dx.doi.org/10.55876/gis8.241206he.

The findings of this study are also based 18,925 dengue sequences available on GISAID up to and accessible at http://dx.doi.org/10.55876/gis8.241206tp. This includes all sequences considered before applying the specified filtering criteria.

The HIV sequences used in this study consist of all available sequences at https://www.hiv.lanl.gov/ before June 2024.

The source code is available under an open source license at https://github.com/INFORM-Africa/AI-viral-lineage-classification.

### Authors’ contribution

JSX, MD, HT, CB, and TdO devised the project. DJvZ developed the models, performed all the experiments, and wrote the paper. MD, JSX, TdO, and CB manage the project and funding. The INFORM Africa research study group supported the project implementation, and all authors read and reviewed it.

## Acknowledgments

We gratefully acknowledge all data contributors, i.e., the Authors and their Originating laboratories responsible for obtaining the specimens, and their Submitting laboratories for generating the genetic sequence and metadata and sharing via the GISAID Initiative, on which this research is based.

The authors also thank P^amela M. Rezende for her help in conceiving the project.

## Additional files

### Additional file 1

Analysis of the feature extraction parameters on the SARS-CoV-2 dataset. (PDF 201 kb)

### Additional file 2

Detailed description of the workings of the selected feature extraction techniques. (PDF 732 kb)

### Additional file 3

Supplemental table acknowledging the authors of the SARS-CoV-2 GISAID sequences. (PDF 21 kb)

### Additional file 4

Supplemental table acknowledging the authors of the dengue GISAID sequences. (PDF 21 kb)

## Footnotes

1 https://github.com/INFORM-Africa/AI-viral-lineage-classification

## References

[1] Zielezinski A, Vinga S, Almeida J, Karlowski WM. Alignment-free sequence comparison: benefits, applications, and tools. Genome Biology. 2017;18(1):186. 10.1186/s13059-017-1319-7.

[2] Karlin S, Altschul S. Methods for assessing the statistical significance of molecular sequence features by using general scoring schemes. Proceedings of the National Academy of Sciences. 1990;87(6):2264–2268. 10.1073/pnas.87.6.2264.

[3] Altschul SF, Gish W, Miller W, Myers EW, Lipman DJ. Basic local alignment search tool. Journal of Molecular Biology. 1990;215(3):403–410. 10.1016/S0022-2836(05)80360-2.

[4] Almeida JS. Sequence analysis by iterated maps, a review. Briefings in Bioinformatics. 2013 10;15(3):369–375. 10.1093/bib/bbt072. https://academic.oup.com/bib/article-pdf/15/3/369/450331/bbt072.pdf.

[5] Lebatteux D, Remita AM, Diallo AB. Toward an Alignment-Free Method for Feature Extraction and Accurate Classification of Viral Sequences. Journal of Computational Biology. 2019;26(6):519–535. PMID: 31050550. 10.1089/cmb.2018.0239. https://doi.org/10.1089/cmb.2018.0239.

[6] Ondov BD, Treangen TJ, Melsted P, Mallonee AB, Bergman NH, Koren S, et al. Mash: fast genome and metagenome distance estimation using MinHash. Genome Biology. 2016 Jun;17(1):132. 10.1186/s13059-016-0997-x.

[7] Kolekar P, Kale M, Kulkarni-Kale U. Alignment-free distance measure based on return time distribution for sequence analysis: applications to clustering, molecular phylogeny and subtyping. Molecular Phylogenetics and Evolution. 2012 Nov;65(2):510–522. 10.1016/j.ympev.2012.07.003.

[8] Leimeister CA, Boden M, Horwege S, Lindner S, Morgenstern B. Fast alignment-free sequence comparison using spaced-word frequencies. Bioinformatics. 2014 04;30(14):1991–1999. 10.1093/bioinformatics/btu177. https://academic.oup.com/bioinformatics/article-pdf/30/14/1991/48925419/bioinformatics_30_14_1991.pdf.

[9] Bonidia RP, Sampaio LDH, Domingues DS, Paschoal AR, Lopes FM, de Carvalho ACPLF, et al. Feature extraction approaches for biological sequences: a comparative study of mathematical features. Briefings in Bioinformatics. 2021 02;22(5):bbab011. 10.1093/bib/bbab011. https://academic.oup.com/bib/article-pdf/22/5/bbab011/40261504/biographies_bbab011.pdf.

[10] Zielezinski A, Girgis HZ, Bernard G, Leimeister CA, Tang K, Dencker T, et al. Benchmarking of alignment-free sequence comparison methods. Genome Biology. 2019;20(1):144. 10.1186/s13059-019-1755-7.

[11] Sims GE, Jun SR, Wu GA, Kim SH. Alignment-free genome comparison with feature frequency profiles (FFP) and optimal resolutions. Proc Natl Acad Sci U S A. 2009 Feb;106(8):2677–2682. 10.1073/pnas.0813249106.

[12] Gao L, Qi J. Whole genome molecular phylogeny of large dsDNA viruses using composition vector method. BMC Evolutionary Biology. 2007;7(1):41. 10.1186/1471-2148-7-41.

[13] Liu ZH, Liu HD, Li JR, Sun X, Jiao D. Base-Base Correlation a Novel Sequence Feature and its Applications. In: 2007 1st International Conference on Bioinformatics and Biomedical Engineering; 2007. p. 370–373.

[14] Hanus P, Dingel J, Zech J, Hagenauer J, Mueller JC. Information Theoretic Distance Measures in Phylogenomics. In: 2007 Information Theory and Applications Workshop; 2007. p. 421–425.

[15] Li M, Badger JH, Chen X, Kwong S, Kearney P, Zhang H. An information-based sequence distance and its application to whole mitochondrial genome phylogeny. Bioinformatics. 2001 02;17(2):149–154. 10.1093/bioinformatics/17.2.149. https://academic.oup.com/bioinformatics/article-pdf/17/2/149/48836819/bioinformatics_17_2_149.pdf.

[16] Otu HH, Sayood K. A new sequence distance measure for phylogenetic tree construction. Bioinformatics. 2003 11;19(16):2122–2130. 10.1093/bioinformatics/btg295. https://academic.oup.com/bioinformatics/article-pdf/19/16/2122/48904654/bioinformatics_19_16_2122.pdf.

[17] Ulitsky I, Burstein D, Tuller T, Chor B. The Average Common Substring Approach to Phylogenomic Reconstruction. Journal of computational biology: a journal of computational molecular cell biology. 2006 04;13:336–50. 10.1089/cmb.2006.13.336.

[18] Leimeister CA, Morgenstern B. kmacs: the k -mismatch average common substring approach to alignment-free sequence comparison. Bioinformatics. 2014 05;30(14):2000–2008. 10.1093/bioinformatics/btu331. https://academic.oup.com/bioinformatics/article-pdf/30/14/2000/48924929/bioinformatics_30_14_2000.pdf.

[19] Haubold B, Pfaffelhuber P, Domazet-Loso M, Wiehe T. Estimating mutation distances from unaligned genomes. Journal of Computational Biology. 2009 Oct;16(10):1487–1500. 10.1089/cmb.2009.0106.

[20] Jeffrey HJ. Chaos game representation of gene structure. Nucleic Acids Res. 1990 Apr;18(8):2163–2170. 10.1093/nar/18.8.2163.

[21] Randhawa GS, Hill KA, Kari L. ML-DSP: Machine Learning with Digital Signal Processing for ultrafast, accurate, and scalable genome classification at all taxonomic levels. BMC Genomics. 2019;20(1):267. 10.1186/s12864-019-5571-y.

[22] Yin R, Luo Z, Kwoh CK. Exploring the Lethality of Human-Adapted Coronavirus Through Alignment-Free Machine Learning Approaches Using Genomic Sequences. Curr Genomics. 2021 Dec;22(8):583–595. 10.2174/1389202923666211221110857.

[23] Storcheus D, Rostamizadeh A, Kumar S. A Survey of Modern Questions and Challenges in Feature Extraction. In: Storcheus D, Rostamizadeh A, Kumar S, editors. Proceedings of the 1st International Workshop on Feature Extraction: Modern Questions and Challenges at NIPS 2015. vol. 44 of Proceedings of Machine Learning Research. Montreal, Canada: PMLR; 2015. p. 1–18. Available from: https://proceedings.mlr.press/v44/storcheus2015survey.html.

[24] Muhammad I, Mukhlash I, Jamhuri M, Iqbal M, Irawan MI. Classification of Covid-19 Variants Using Boosting Algorithm. In: 2022 9th International Conference on Electrical Engineering, Computer Science and Informatics (EECSI); 2022. p. 29–34.

[25] Lebatteux D, Soudeyns H, Boucoiran I, Gantt S, Diallo A. Machine learning-based approach KEVOLVE efficiently identifies SARS-CoV-2 variant-specific genomic signatures. PLoS One. 2024;19(1):e0296627. 10.1371/journal.pone.0296627.

[26] Cacciabue M, Aguilera P, Gismondi M, Taboga O. Covidex: An ultrafast and accurate tool for SARS-CoV-2 subtyping. Infection, Genetics and Evolution. 2022 April;99:105261. Epub 2022 Feb 26. 10.1016/j.meegid.2022.105261.

[27] Khare S, Gurry C, Freitas L, Schultz M, Bach G, Diallo A, et al. GISAID’s Role in Pandemic Response. China CDC Weekly. 2021 December;3(49):1049–1051. 10.46234/ccdcw2021.255.

[28] Apetrei C, Hahn B, Rambaut A, Wolinsky S, Brister JR, Keele B, et al., editors. HIV Sequence Compendium 2021. Los Alamos, NM: Theoretical Biology and Biophysics Group, Los Alamos National Laboratory; 2021. LA-UR-23-22840.

[29] O’Toole A, Pybus OG, Abram ME, Kelly EJ, Rambaut A. Pango lineage designation and assignment using SARS-CoV-2 spike gene nucleotide sequences. BMC Genomics. 2022 Feb;23(1):121. 10.1186/s12864-022-08358-2.

