## Supplementary material for "Alignment-Free Viral Sequence Classification at Scale": Analysis of the feature extraction parameters on the SARS-CoV-2 dataset.

### Additional file 1

#### Analysis of Feature Extraction Parameters

We compared the effects of removing versus randomly replacing degenerate nucleotides across all AF methods. Figure 1 shows a comparison of these two strategies for the SARS-CoV-2 dataset.

For all techniques except GSP, the Macro F1 performance of the Random Forest models did not differ significantly between removing and replacing degenerate nucleotides. In word-based models, removing degenerate nucleotides proved to be marginally superior, while for MASH, the opposite was true.

GSP demonstrated a greater difference between the two strategies, with the replacement of degenerate nucleotides proving more optimal. This is likely because GSP relies more on the positioning of nucleotides compared to other techniques. By replacing nucleotides instead of removing them, the relative positions of the nucleotides remain intact, thereby preserving the positional information.

We also provide a comparison of the effect of increasing the value of $k$, the length of the word for the word-based methods, in Figure 2. For FCGR, the word size $k$ corresponds to the resolution of the generated images.

All four word-based models exhibit a similar trajectory in terms of Macro F1 classification performance with increasing word length. In all cases, increasing the length of the word improved Macro F1 performance, with diminishing returns as the length increased. Once the word length reaches seven, we observe only marginal improvements. The anticipated improvement with increasing word length must be weighed against the increased computational complexity. Increasing the word length by a single character quadruples the number of features for word-based methods, significantly impacting computational requirements.

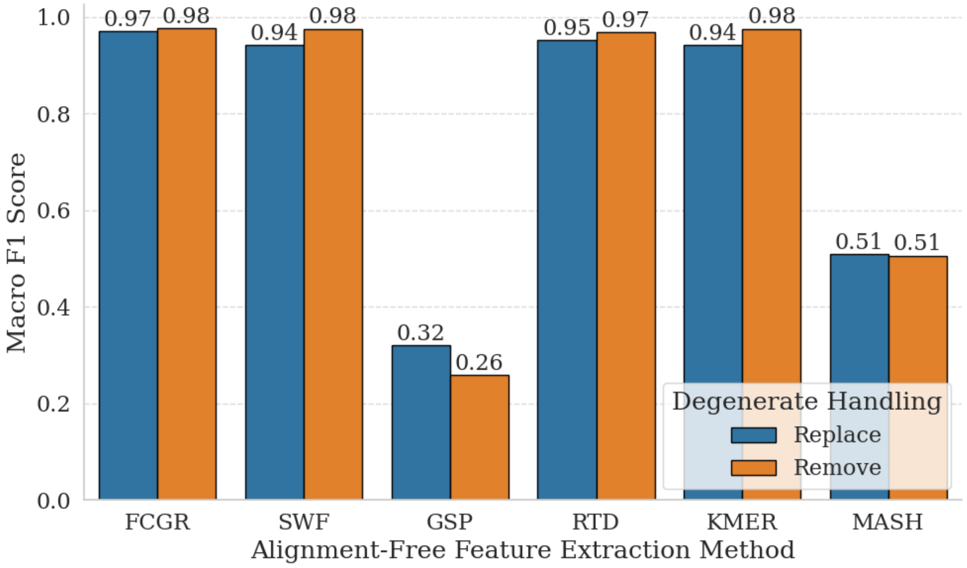


**Fig. 1** A comparison of Macro F1 classification performance on the SARS-CoV-2 test set for different AF feature extraction methods when replacing (Blue) vs removing (Orange) degenerate/ambiguous nucleotide bases.

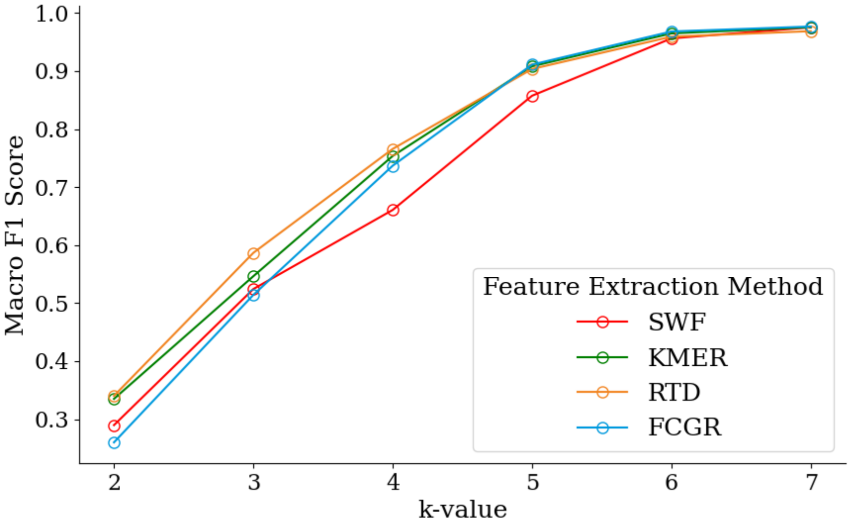


**Fig. 2** A comparison of Macro F1 classification performance on the SARS-CoV-2 test set for word-based AF feature extraction methods when using different word sizes $k$. In the case of FCGR, the word size corresponds to a resolution of $2^k$.
