## Supplementary material for "Alignment-Free Viral Sequence Classification at Scale": Detailed description of the workings of the selected feature extraction techniques.

### Additional file 2

### Alignment-Free Sequence Comparison Methods

#### *k*-mer Count

$k$-mer counting deconstructs genomic sequences into all conceivable overlapping subsequences, also known as words, of a specific length $k$ [1, 2]. Such overlapping words are termed "$k$-mers", with lenght $k$. In genomic sequences, with an alphabet comprising A, C, T, G, the total potential $k$-mers for each $k$ value is $4^k$. To construct a feature vector, one counts the number of occurrences of each possible $k$-mer. In particular, to distinguish between highly similar sequences, larger values $k$ are typically required, although at the expense of increased computational costs [3]. Figure 1 depicts an illustrative example of the $k$-mer construction process.

#### Return Time Distribution (RTD)

Return Time Distribution (RTD) [4] approaches $k$-mer occurrence through the lens of recurrence intervals rather than direct frequency. RTD calculates the mean ($\mu$) and standard deviation ($\sigma$) of the intervals at which each $k$-mer reappears, thus characterizing each sequence with a numerical vector twice the size of the total number of potential $k$-mers.

#### Spaced Word Frequencies

Spaced Word Frequencies [2] leverage a nuanced approach within word frequency analysis by employing patterns that intersperse "match" positions, which requires exact character correspondence, with "don't care" positions, where character identity is irrelevant. This methodology facilitates the identification of "spaced words" within sequences, focusing solely on the "care" positions, thus allowing for the accommodation of sequence variations or mutations [2], and diminishing the statistical dependence

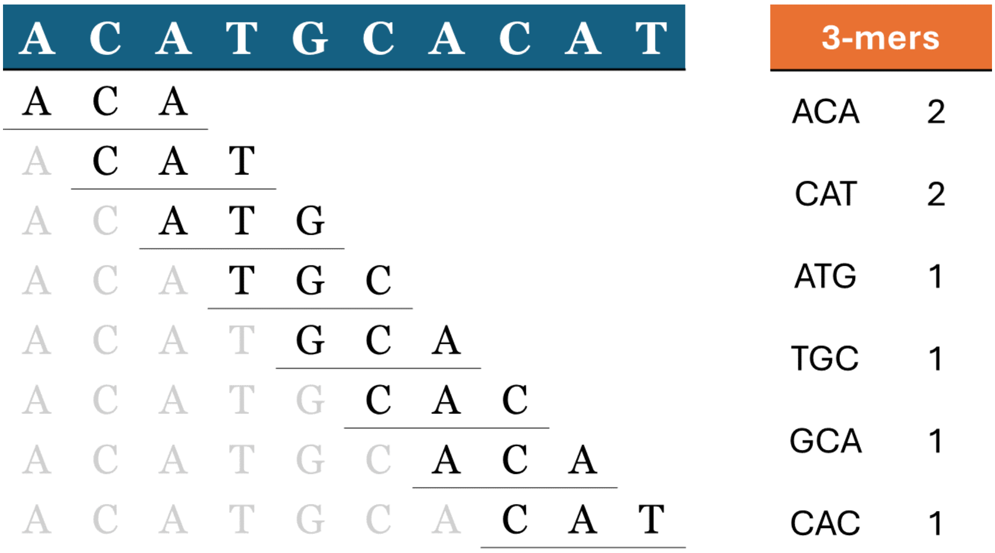


**Fig. 1** Illustrative example of the $k$-mer counting procedure for the sequence "ACATGCACAT" with $k = 3$. In this example, only $k$-mers present in the sequence are shown, however, in constructing a full feature vector, zero counts for all other possible 3-mers must also be included.

observed in adjacent word matches prevalent in contiguous word analysis [2, 3]. Figure 2 demonstrates the use of spaced words for an illustrative example.

#### Mash

Mash [5] builds upon the MinHash technique, traditionally used for identifying nearly identical web pages and images, and adapting it to genomic data sets. MinHash is used to efficiently estimate the similarity between large sets by hashing and comparing subsets of their elements. A hash function converts input data such as a character sequences into fixed-length numerical values to allow for easier indexing and comparisons. In the case of Mash, $k$-mers are extracted, hashed, and sorted, with the $s$ smallest hash values considered the Mash 'sketch'. Figure 3 shows an illustration of Mash sketch construction.

The Jaccard index is a similarity measure between two sets, calculated by dividing the size of the intersection of the sets by the size of their union. Mash sketches can therefore be used to estimate the Jaccard index between $k$-mer sets from two sequences by calculating the Jaccard index for the elements of the two sketches only. The final Mash distance, modelled under a Poisson process, is calculated using the relationship between the expected fraction of unchanged $k$-mers and the Jaccard estimate.

#### Chaos Game Representation

The Chaos Game Representation (CGR) [6], introduced by Jeffrey in 1990, is a method for identifying patterns in DNA sequences using principles from chaos theory. The

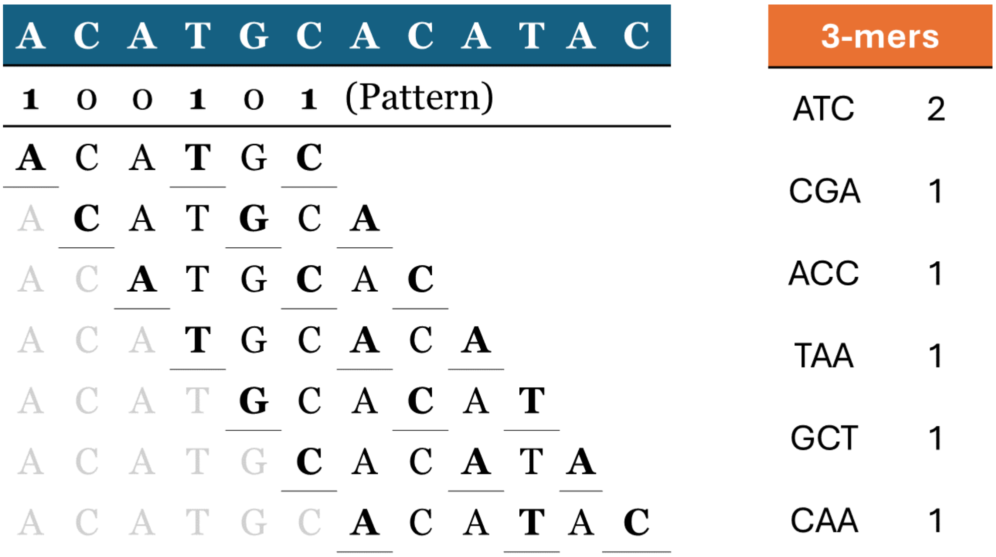


**Fig. 2** Demonstration of spaced word calculation for the sequence "ACATGCACATAC". In this example, the selected pattern "100101" has a weight of three for the three match positions. During the feature construction, the pattern is shifted along the sequence with single increments. At each position, $k$-mers are constructed considering only the match positions (1's).

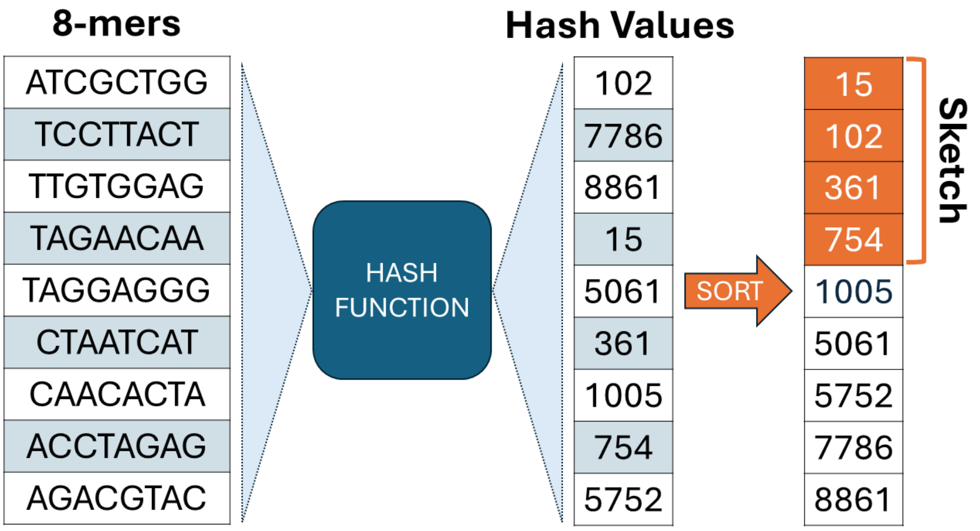


**Fig. 3** To construct a Mash sketch, each $k$-mer is hashed to a fixed-size numerical value using a hash function. Mash selects the smallest hashes, the bottom-sketch, based on a predefined sketch size $s$, which is four in this example.

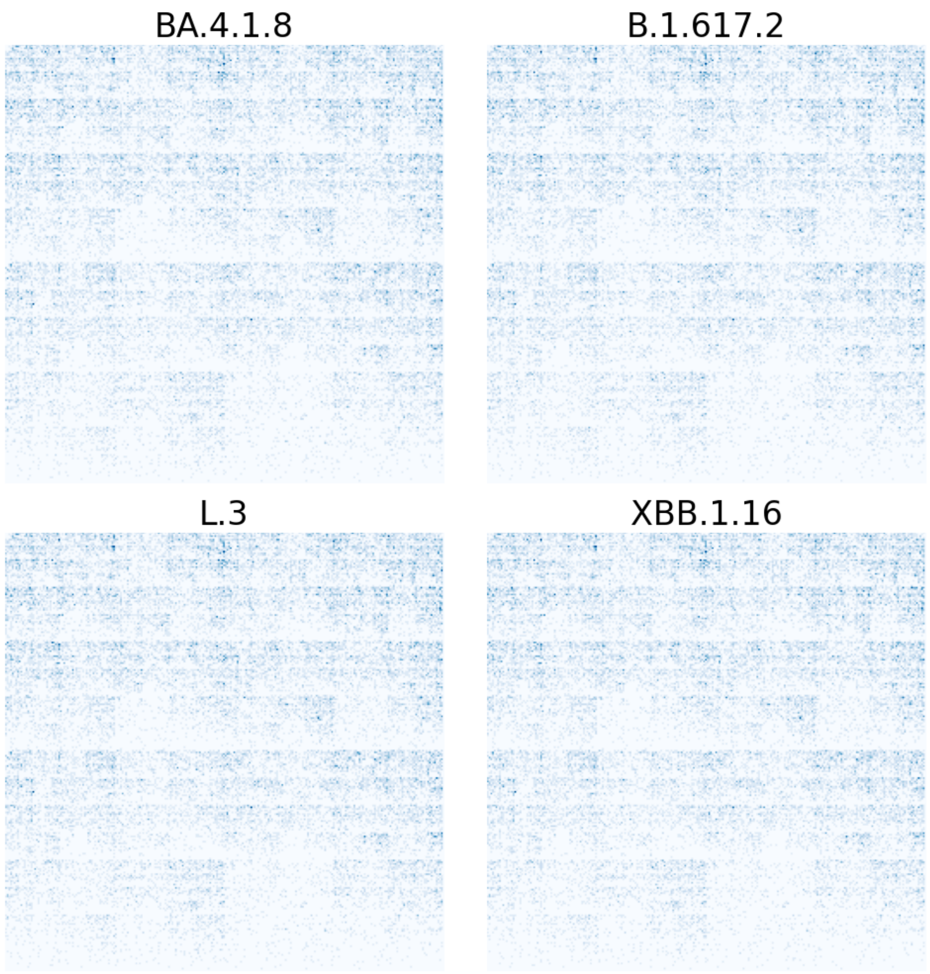


**Fig. 4** An example of Frequency Chaos Game Representations for four distinct SARS-CoV-2 lineages.

method generates a visual pattern of dots within a unit square, wherein each corner represents one of the DNA bases: A, C, G, and T. Starting in the center of the unit square, and reading the first nucleotide, points are plotted by moving halfway towards the corner corresponding to each nucleotide in the sequence. This process continues, creating a pattern that captures a visual fingerprint for each sequence. The resulting image can be described as a vector of chronological co-ordinate pairs, but also by considering the frequency of dots within grid squares. Figure 4 provides example visualizations of Frequency Chaos Game Representation (FCGR) images.

#### Genomic Signal Processing

Genomic Signal Processing (GSP) integrates Digital Signal Processing (DSP) techniques with genomic data, using predefined numeric mappings to first convert DNA sequences into discrete numerical sequences [7, 8]. Table 1 contains some of the most popular numeric mapping schemes [9].

**Table 1** Encoding Schemes for the Sequence CGAT

| **Representation** | **Substitutions: [A, C, G, T]** |
|---|---|
| Integer | [2, 1, 3, 0] |
| Real | [1.5, 0.5, -0.5, -1.5] |
| Atomic | [70, 58, 78, 6] |
| EIIP | [0.1260, 0.1340, 0.0806, 0.1335] |
| PP | [-1, 1, -1, 1] |
| Just-A | [1, 0, 0, 0] |
| Just-C | [0, 1, 0, 0] |
| Just-G | [0, 0, 1, 0] |
| Just-T | [0, 0, 0, 1] |

Following a length normalization procedure, each numeric nucleotide sequence is mapped to a complex value through the Discrete Fourier Transform (DFT) function $f(\cdot)$.

$$F_i(k) = \sum_{j=0}^{l-1} f(S_i(j)) \cdot e^{-\frac{2\pi i k j}{l}}$$

The transformed signal for each sequence is represented as a vector $F_i$. Here, $F_i(k)$ is the Fourier coefficient at frequency $k$ for the sequence $S_i$. The term $f(S_i(j))$ denotes the numerical value of the $j$-th nucleotide after mapping. After computing the DFT, we obtain the magnitude spectrum $M_i$, the absolute value of $F_i$. Once these spectrums are obtained, the distance between two sequences can be computed through the use of Pearson correlation dissimilarity.

### References

[1] Manekar SC, Sathe SR. A benchmark study of k-mer counting methods for high-throughput sequencing. GigaScience. 2018 10;7(12):giy125. https://doi.org/10.1093/gigascience/giy125. https://academic.oup.com/gigascience/article-pdf/7/12/giy125/27011542/giy125_reviewer_3_report_(original_submission).pdf.

[2] Leimeister CA, Boden M, Horwege S, Lindner S, Morgenstern B. Fast alignment-free sequence comparison using spaced-word frequencies. Bioinformatics. 2014 04;30(14):1991–1999. https://doi.org/10.1093/bioinformatics/btu177. https://academic.oup.com/bioinformatics/article-pdf/30/14/1991/48925419/bioinformatics_30_14_1991.pdf.

[3] Zielezinski A, Vinga S, Almeida J, Karlowski WM. Alignment-free sequence comparison: benefits, applications, and tools. Genome Biology. 2017;18(1):186. https://doi.org/10.1186/s13059-017-1319-7.

[4] Kolekar P, Kale M, Kulkarni-Kale U. Alignment-free distance measure based on return time distribution for sequence analysis: applications to clustering, molecular phylogeny and subtyping. Molecular Phylogenetics and Evolution. 2012 Nov;65(2):510–522. https://doi.org/10.1016/j.ympev.2012.07.003.

[5] Ondov BD, Treangen TJ, Melsted P, Mallonee AB, Bergman NH, Koren S, et al. Mash: fast genome and metagenome distance estimation using MinHash. Genome Biology. 2016 Jun;17(1):132. https://doi.org/10.1186/s13059-016-0997-x.

[6] Jeffrey HJ. Chaos game representation of gene structure. Nucleic Acids Res. 1990 Apr;18(8):2163–2170. https://doi.org/10.1093/nar/18.8.2163.

[7] Randhawa GS, Hill KA, Kari L. ML-DSP: Machine Learning with Digital Signal Processing for ultrafast, accurate, and scalable genome classification at all taxonomic levels. BMC Genomics. 2019;20(1):267. https://doi.org/10.1186/s12864-019-5571-y.

[8] Yin R, Luo Z, Kwoh CK. Exploring the Lethality of Human-Adapted Coronavirus Through Alignment-Free Machine Learning Approaches Using Genomic Sequences. Curr Genomics. 2021 Dec;22(8):583–595. https://doi.org/10.2174/1389202923666211221110857.

[9] Kwan HK, Arniker SB. Numerical representation of DNA sequences. In: 2009 IEEE International Conference on Electro/Information Technology; 2009. p. 307–310.
