## Supplemental table acknowledging the authors of the SARS-CoV-2 GISAID sequences. for "Alignment-Free Viral Sequence Classification at Scale"

**Data Availability**

GISAID Identifier: EPI_SET_241206he
doi: 10.55876/gis8.241206he

All genome sequences and associated metadata in this dataset are published in GISAID's EpiCoV database. To view the contributors of each individual sequence with details such as accession number, Virus name, Collection date, Originating Lab and Submitting Lab and the list of Authors, visit 10.55876/gis8.241206he

**Data Snapshot**

- EPI_SET_241206he is composed of 297,184 individual genome sequences.
- The collection dates range from 2019-12-31 to 2024-04-15;
- Data were collected in 206 countries and territories;
- All sequences in this dataset are compared relative to hCoV-19/Wuhan/WIV04/2019 (WIV04), the official reference sequence employed by GISAID (EPI_ISL_402124). Learn more at https://gisaid.org/WIV04.
