## Supplemental table acknowledging the authors of the dengue GISAID sequences. for "Alignment-Free Viral Sequence Classification at Scale"

### Data Availability


GISAID Identifier: EPI_SET_241206tp
doi: 10.55876/gis8.241206tp


All genome sequences and associated metadata in this dataset are published in GISAID’s EpiArbo database. To view the contributors of each individual sequence with details such as accession number, Virus name, Collection date, Originating Lab and Submitting Lab and the list of Authors, visit 10.55876/gis8.241206tp

### Data Snapshot

- EPI_SET_241206tp is composed of 18,925 individual genome sequences.
- The collection dates range from 1944-01-01 to 2024-05-02;
- Data were collected in 111 countries and territories;
- All sequences in this dataset are compared relative to the official reference sequence employed by GISAID.
